# An Analysis of Characterized Plant Sesquiterpene Synthases

**DOI:** 10.1101/466839

**Authors:** Janani Durairaj, Alice Di Girolamo, Harro J Bouwmeester, Dick de Ridder, Jules Beekwilder, Aalt DJ van Dijk

## Abstract

Plants exhibit a vast array of sesquiterpenes, C15 hydrocarbons which often function as herbivore-repellents or pollinator-attractants. These in turn are produced by a diverse range of sesquiterpene synthases. A comprehensive analysis of these enzymes in terms of product specificity has been hampered by the lack of a centralized resource of sufficient functionally annotated sequence data. To address this, we have gathered 262 plant sesquiterpene synthase sequences with experimentally characterized products. The annotated enzyme sequences allowed for an analysis of terpene synthase motifs, leading to the extension of one motif and recognition of a variant of another. In addition, putative terpene synthase sequences were obtained from various resources and compared with the annotated sesquiterpene synthases. This analysis indicated regions of terpene synthase sequence space which so far are unexplored experimentally. Finally, we present a case describing mutational studies on residues altering product specificity, for which we analyzed conservation in our database. This demonstrates an application of our database in choosing likely-functional residues for mutagenesis studies aimed at understanding or changing sesquiterpene synthase product specificity.

## 1. Introduction

The terpenome represents a huge, ancient and diverse family of natural products. In addition to terpenes, it also encompasses steroids and carotenoids, comprising more than 60,000 members [1]. These compounds all derive from the same 5-carbon precursor units, coupled together linearly and then cyclized, rearranged, and modified in various ways. Terpenes serve many roles in plants, for example as toxins against herbivores or pathogens, or as attractants for pollinators [2]. In turn, terpenes extracted from plants are used by mankind for a range of applications - as pharmaceutical agents, insecticides, preservatives, fragrances, and flavors [3].

Terpenes are built from 5-carbon isoprenoid units and they mainly exist as monoterpenes (C10), sesquiterpenes (C15) or diterpenes (C20), based on the number of such units used. In each case, a linear substrate loses a diphosphate group, usually cyclizes and then undergoes a variety of carbocation rearrangements. Though the exact number of sesquiterpenes found in nature is hard to determine, Tian *et al* estimated computationally that the number of sesquiterpene intermediates far outnumber those of monoterpenes, due to the increase in chain length [4]. Interestingly, sesquiterpenes found in nature can be divided into seven groups based on their parent cation and the first cyclization step in their formation [5]. Hence the extreme diversity of chemical compounds with desirable fragrances or medicinal properties is based on just seven initial carbocations. This makes the enzymes catalyzing their formation both interesting and difficult to characterize functionally.

Each plant species is capable of synthesizing a number of sesquiterpenes using a specialized class of enzymes called sesquiterpene synthases (STSs). First, a farnesyl diphosphate synthase, produces the C15 substrate for STSs, farnesyl diphosphate (FPP), from the C5-unit isopentenyl diphosphate (IPP) and its isomer dimethylallyl diphosphate (DMAPP) [6]. STSs then create the myriad of sesquiterpenes found in nature by catalyzing carbocation formation from the linear FPP followed by a series of cyclizations and rearrangements (Figure 1). Products are formed from intermediate carbocations after deprotonation, phosphorylation, or hydration [4].

**Figure 1:**
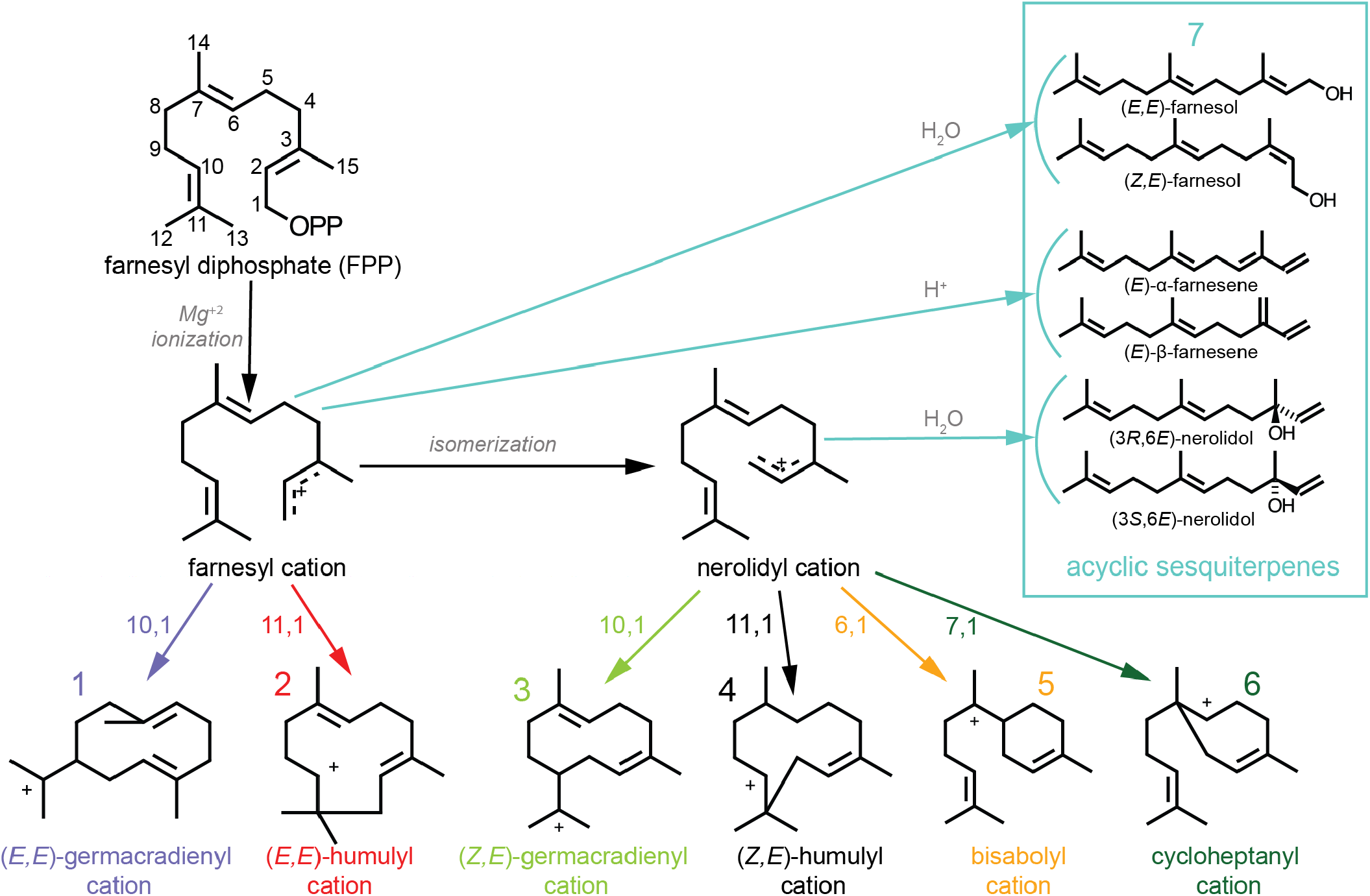
The reaction mechanism of sesquiterpene production starts with farnesyl diphosphate (FPP). Loss of the diphosphate moiety (OPP) leads to farnesyl cation formation. The farnesyl cation can subsequently be converted to the nerolidyl cation. Possible cyclizations for both cations are indicated in the figure. The subsequently formed cyclic cations undergo further modifications and rearrangements to form sesquiterpenes. An alternative route is to form acyclic sesquiterpenes from either the farnesyl or the nerolidyl cation. These different product-precursors are used to classify the different sesquiterpenes and their synthases.

The STSs themselves represent a very diverse set of enzymes with a wide range of sequence similarities, despite having a common structural fold shared by plant, animal, fungal, and bacterial terpene synthases (TPSs) [7]. Hence, prediction of enzyme function from sequence is highly challenging in the case of STSs. Moreover, sequence diversity in STSs is not dependant on the products formed. This problem has been addressed so far by inspection of TPS structures [7] and by mutational analyses that attempt to change the product of a synthase with the smallest number of residue changes [8]. The former, though an attractive approach, is limited especially in plants due to the sparsity of experimentally determined structures, while the latter often leads to unnatural enzymes with lower catalytic activity than their wild-type parents. Characterization of multiple TPSs from the same species by the same study has allowed for some small-scale sequence comparison of those synthases [9, 10]. However, no previous attempts have been made to compare all experimentally characterized plant STS sequences according to the products that they form. We have collated a curated database of plant STSs with characterized products from literature. This database can be accessed at www.bioinformatics.nl/sesquiterpene/synthasedb.

With this database and aforementioned product grouping scheme, the active domain sequences of 262 plant STSs were analyzed in terms of the precursor carbocations of their products. These were also compared with the many yet-uncharacterized putative TPS enzymes. Residues from previous product-changing mutational studies were mapped on our database of enzymes, indicating conservation of the corresponding positions across groups of sequences forming different product cations. This demonstrates the usefulness of our database in finding residues involved in STS product specificity.

## 2. Results and Discussion

### 2.1. Database of characterized STSs

To obtain a comprehensive set of annotated STSs, our starting point was the SwissProt database, a subset of UniProt [11] in which a curated and annotated set of proteins is available. This provided a set of 104 STSs. In addition, we manually reviewed literature linked to enzymes with the characteristic TPS domain in TremBl, the uncurated subset of UniProt. In this way, the number of curated plant STS sequences with experimentally characterized product data in the database was more than doubled.

We present a database of 262 manually curated characterized plant STSs, shown in Table 1. The enzymes originate from a hundred different plant species and collectively account for the production of 117 different sesquiterpenes. Such a large number of possible products makes it difficult to find enough enzymes with the same product for a meaningful analysis of product specificity. To solve this, the sequences were divided into seven groups, making use of the sesquiterpene precursor carbocation scheme as specified by Degenhardt *et al*. [5], described in Figure 1. The reaction cascade of an STS is initiated by metal-mediated removal of the diphosphate anion in the FPP substrate, leading to the formation of a transoid (2*E*,6*E*)-farnesyl cation (farnesyl cation) which, due to its constrained structure, can undergo cyclization only at its C10-C11 double bond leading to cations 1 or 2 in Figure 1. However, the farnesyl cation can also isomerize to form a cisoid (2*Z*,6*E*)-farnesyl cation (nerolidyl cation). The nerolidyl cation, in addition to a C1-attack on the C10-C11 double bond to form cations 3 or 4, can also undergo cyclization at its C6-C7 double bond, forming cations 5 or 6. These carbocations undergo multiple further skeletal rearrangements, cyclizations, hydride or methyl shifts, and other modifications to form the final products of the enzyme [5]. Along with this myriad of cyclic products, acyclic sesquiterpenes can also be formed from either the farnesyl or the nerolidyl cation through proton loss or addition of water [5, 12, 13]. This schematic of carbocations derived from FPP can be used to divide sesquiterpenes produced by plants into seven groups - both based on their parent cation (farnesyl or nerolidyl) and the first cyclization that occurs (by attack of the carbocation on the 10,1-; 11,1-; 6,1-; or 7,1- double bond; or acyclic). For an STS enzyme, the carbocation of its major product is then used to determine its group in Table 1.

**Table 1.** Number of characterized plant STS sequences, species, and products covered in each product group. (-)-germacrene D synthases are shown separately as discussed in the text. 1=Angiosperms, 2=Gymnosperms, 3=Nonseed

| Major Product Group | Cation / Cyclization | Number of sequences |  |  |  | Number of species |  |  |  | Number of products |
| --- | --- | --- | --- | --- | --- | --- | --- | --- | --- | --- |
|  |  | A<sup>1</sup> | G<sup>2</sup> | N<sup>3</sup> | Total | A<sup>1</sup> | G<sup>2</sup> | N<sup>3</sup> | Total |  |
| 1 | 10,1 / farnesyl | 77 | 1 | 3 | 81 | 44 | 1 | 3 | 48 | 43 |
| 2 | 11,1 / farnesyl | 42 | 3 | 3 | 48 | 32 | 3 | 3 | 38 | 11 |
| 3 | 10,1 / nerolidyl | 19 | 1 | 1 | 21 | 16 | 1 | 1 | 18 | 20 |
| 4 | 11,1 / nerolidyl | 0 | 4 | 0 | 4 | 0 | 4 | 0 | 4 | 3 |
| 5 | 6,1 / nerolidyl | 44 | 3 | 2 | 49 | 23 | 3 | 2 | 28 | 32 |
| 6 | 7,1 / nerolidyl | 0 | 0 | 1 | 1 | 0 | 0 | 1 | 1 | 1 |
| 7 | acyclic | 43 | 4 | 3 | 50 | 23 | 4 | 3 | 30 | 6 |
| - | (-)-germacrene D | 8 | 0 | 0 | 8 | 6 | 0 | 0 | 6 | 1 |
| Total |  | 233 | 16 | 13 | 262 | 84 | 8 | 9 | 101 | 117 |

This division of STSs is in general straightforward even when multiple products are formed by one enzyme. Specifically, of the 98 sequences which also have minor products (Supplementary Table T1), only 17 have minor products whose precursor carbocation differs from the major product’s. Nine of these produce acyclic products in addition to their major product. This could be the result of incomplete cyclization caused by premature termination of intermediates [14]. Eight enzymes in the database either produce (-)-germacrene D or they produce germacrene D and the chirality was not determined during the enzyme’s characterization. (-)-germacrene D can be formed via a 10,1- or a 11,1- cyclization of the farnesyl cation (cation 1 or 2). Though each enzyme is likely to only follow one cyclization route to form its product, this route has so far not been determined, so these sequences are shown separately in Table 1 and in the remainder of the text. The existence of other sesquiterpenes which can be formed via different cyclization routes cannot be ruled out, however in our analysis we stick to the cyclization routes provided by IUBMB’s *Enzyme Nomenclature* Supplement 24 (2018) [15] in order to determine the precursor carbocation for a given sesquiterpene.

The database contains 233 angiosperm STSs, 16 gymnosperm enzymes from coniferous species and 13 enzymes from nonseed plants such as mosses and ferns. As described by Chen *et al*. [16], the latter species have TPSs which are more related to microbial TPSs than those from spermatophytes. Information on each of the 262 enzymes, including the sequence, species, Uniprot ID, products (major and minor), product type, and Pubmed ID of the paper detailing its experimental characterization, is available as a web service at www.bioinformatics.nl/sesquiterpene/synthasedb. The service supports searching, sorting and downloading of all or subsets of the data.

On average, the enzymes comprise of 553 ± 56 residues. The tertiary structure of STS enzymes usually comprises of two alpha-helical domains [17]. The N-terminal domain is considered relictual in plant STSs, while the C-terminal domain, consisting of an α-helical bundle, is catalytically active [18, 7]. The hydrophobic active site pocket in this domain is formed by six α-helices, closed by two loops. Supplementary Table T2 gives a list of plant STS structures from the Protein Data Bank (PDB) [19]. The C-terminal sub-sequences containing the active site are obtained from each enzyme in the database using information from Pfam [20], and consist of 266 ± 7 residues. Since product and intermediate formation occurs in the active site pocket, from this point on we concentrate on the C-terminal subsequences of TPSs. Supplementary Figure S1 shows the pairwise sequence identity scores for each pair of C-terminal domain sub-sequences for the enzymes in the database, hierarchically clustered and coloured by product cation type. It can be seen that many pairs of sequences have less than 40% sequence identity.

The phylogenetic tree of C-terminal sub-sequences of all 262 enzymes (Figure 2) shows some grouping of spermatophyte enzymes based on their product precursor. In general, the neighbor of an enzyme is from the same or related species, and if there are enough examples from the same species then some product-based grouping is seen. For example, the clades containing mostly enzymes from *Zea mays* on the right are separated based on the product carbocation of the enzyme even while being grouped by the species. However, this is not a consistent trend - enzymes from *Vitis* and *Santalum* at the top of the tree group mainly by species and not by product type. In fact, the three *Santalum* synthase sequences marked in Figure 2, making products derived from three different cyclic carbocations, have more than 90% in common. In any case, the product group of an enzyme from a species not present in the tree is nearly impossible to predict, while enzymes from species which are less represented in the tree can also be difficult to classify. In addition, clades forming predominantly one product carbocation are seen in many different parts of the tree, showing that strongly varying sequences can catalyze the same cyclization reaction and even produce the same product, such as the two marked *β*-caryophyllene synthases from *Arabidposis lyrata* and *Zea perennis* which have a sequence identity less than 30%. Hence phylogenetic analysis is biased and cannot be an accurate predictor of TPS product specificity.

**Figure 2:**
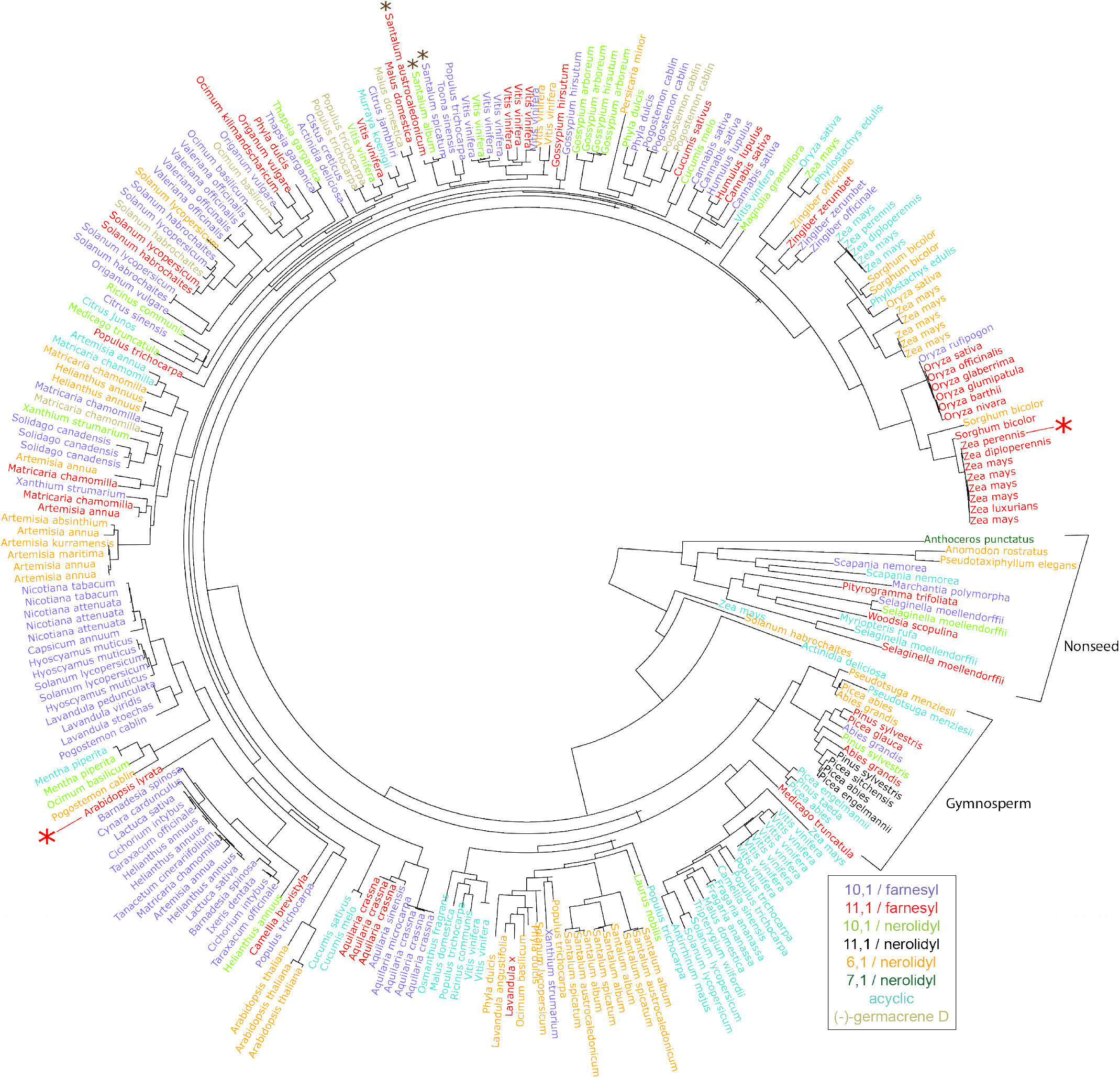
Phylogenetic tree of C-terminal sub-sequences for characterized plant STSs, coloured according to the major product’s initial carbocation (see Figure 1). Nonseed and gymnosperm clades are indicated separately. Red and brown asterisks mark cases discussed in the text: red - two β-caryophyllene synthases from *Arabidopsis lyrata* and *Zea perennis* which have less than 30% pairwise sequence identity; brown - three synthases from *Santalum* with higher than 90% sequence identity.

The clade containing all the nonseed plant STSs is clearly separate from the spermatophyte sequences. The enzyme from *Anthoceros punctatus*, a bryophyte, is the only sequence in the database producing a 7,1/nerolidyl-derived product (*β*-acoradiene) and is hence an out-group both in terms of species as well as product carbocation. Comparing nonseed plant sequences to the more typical plant TPS sequences would be futile, both due to their homology with microbial enzymes and their low numbers in the database, hence they are excluded from the remainder of the analysis.

### 2.2. Chemical similarities between sesquiterpenes

Each of the seven possible sesquiterpene precursors (Figure 1) usually undergoes a wide range of further rearrangements, cyclizations, and modifications, catalyzed by the STS enzyme, to finally result in a sesquiterpene product. To start exploring the enzyme grouping scheme, we initially investigated whether similarities between the final sesquiterpene chemical structures would reflect the parent carbocations involved in their production. To this end, chemical similarities between sesquiterpenes with the same parent cation were compared to similarities between those without. Chemical similarities were measured using Dice similarity [21] between extended connectivity fingerprints, as described by Rogers *et al*. [22]. Similarities between 165 sesquiterpenes are plotted using multi-dimensional scaling (MDS), in Figure 3*a*, with the color representative of the precursor cation. These 165 compounds collectively represent every enantiomer of the 117 sesquiterpenes produced by the enzymes in our database, since many of the experimental characterization studies used to build the database did not resolve the chirality of the STS’s product. MDS is a technique used to visualize the level of similarity of individual objects in a dataset using a distance matrix, such that the between-object distances are preserved as well as possible. Therefore, two objects appearing close to each other in the MDS plot represent sesquiterpenes which likely have a high chemical similarity, while those further away have lower similarity. Acyclic sesquiterpenes are clearly distinguishable in the plot, as they are linear in nature. Interestingly, many products derived from the 6,1-cyclized cation (cation 5) are also distinct from those derived from 10,1- or 11,1-cyclized cations despite further cyclizations and rearrangements after this first step. They cluster midway between the acyclic and other cyclic products, which makes sense given the presence of an acyclic tail portion in cation 5. The sesquiterpenes formed from the other cyclic cations seem less distinguishable.

**Figure 3:**
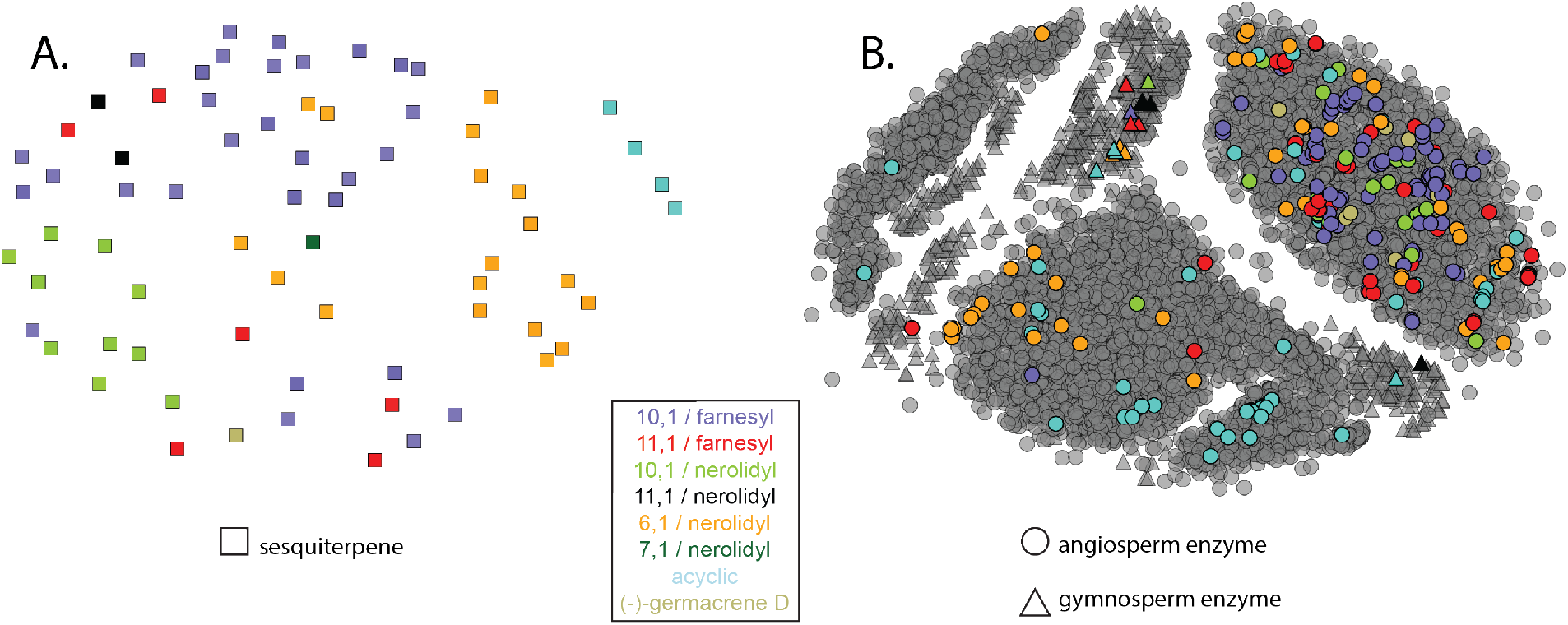
A. MDS plot of 165 sesquiterpenes found in nature, based on chemical fingerprint similarities. Each square represents a sesquiterpene and the more chemically similar two sesquiterpenes are, the closer they are placed in the plot. Colours are based on the sesquiterpene’s precursor carbocation. B. MDS plot of TPS C-terminal domain sub-sequences with coloring based on STS major product carbocation. Unknown proteins which are likely to be TPSs are shown in gray. The more similar two sequences are, the closer they are in the plot.

### 2.3. Characterized sequence space

Though a manual literature search gave us access to more functionally characterized TPS sequences, there is a large and steadily growing number of protein sequences present in various databases which have not been characterized at all. Many of these proteins are potential TPSs which contain the characteristic, catalytic site containing, C-terminal domain. Comparing uncharacterized and characterized enzymes may give indications of the nature of an uncharacterized enzyme, in particular about the cyclization route it is likely to take, thereby assisting in the setup of experiments for functional characterization.

To explore this, an MDS plot was made of C-terminal sub-sequences of the 249 spermatophyte enzymes in our database with those of 6278 other spermatophyte TPS-like sequences, obtained from sequenced genomes and transcriptomes. These 6278 sequences are, to the best of our knowledge, uncharacterized. Figure 3*b* shows this plot where the colors represent the product precursor carbocation of characterized STSs and the uncharacterized sequences are shown in gray. Similar sequences are depicted closer together in the plot.

Figure 3*b* has a few commonalities with the MDS plot of chemical similarities between sesquiterpenes, Figure 3*a*. Many sequences catalyzing acyclic products as well as those derived from cation 5 cluster separately from the others. In fact, the enzymes making nerolidol, an acyclic sesquiterpene, cluster separately at the bottom right of the plot (light blue), leading us to hypothesize that perhaps many of the other uncharacterized STSs in this area also catalyze the formation of nerolidol. A second similarity is that enzymes forming products derived from 10,1- and 11,1- cyclized cations are difficult to distinguish. This again confirms, as was seen in the phylogenetic tree (Figure 2), that overall sequence similarity by itself cannot be an accurate guide to product specificity.

The uncharacterized sequences depicted in Figure 3*b* could be mono-, di-, or sesquiterpene synthases. Supplementary Figure S2 shows 57 monoterpene synthases and 20 diterpene synthases from SwissProt, along with the 249 STSs in our database. Despite the skewed numbers, a separation between mono- and sesquiterpene synthases can be seen, indicating areas of the sequence space where more STSs are likely to be found.

Product specificity is even harder to identify in the case of gymnosperm synthases, as insufficient data is available to separate enzymes with different product cations. It has been noted before that gymnosperm TPSs resemble each other more than they do their angiosperm counterparts, regardless of catalytic activity [23, 24]. The enzymes from these species may be more informative if analyzed separately but this would require more gymnosperm sequences to be functionally annotated.

### 2.4. Comparing known TPS motifs across sequences

A database such as ours allows for a comparison of residues in previously studied structural elements across many STS sequences. A thorough study of TPS structures has led to the identification of several motifs important for catalytic activity [7]. In the case of STSs, the hydrophobic moiety of the STS substrate, FPP, is directed into the active site cavity, to undergo the cyclizations and rearrangements described in Figure 1. Studies on STS structures have proposed that the diphosphate moiety is captured by the motif RxR and divalent metal ions like *Mg*^+2^ or *Mn*^+2^, which are themselves bound by motifs DDxxD and NSE/DTE, at the entrance of the active site [25]. Here, we compare these three motifs across the sequences in our database. Figure 4*a* shows the motifs discussed below on a tobacco aristolochene synthase structure [25]. Figure 4*b* shows each motif on a schematic representation of the alignment of all C-terminal sub-sequences in the database.

**Figure 4:**
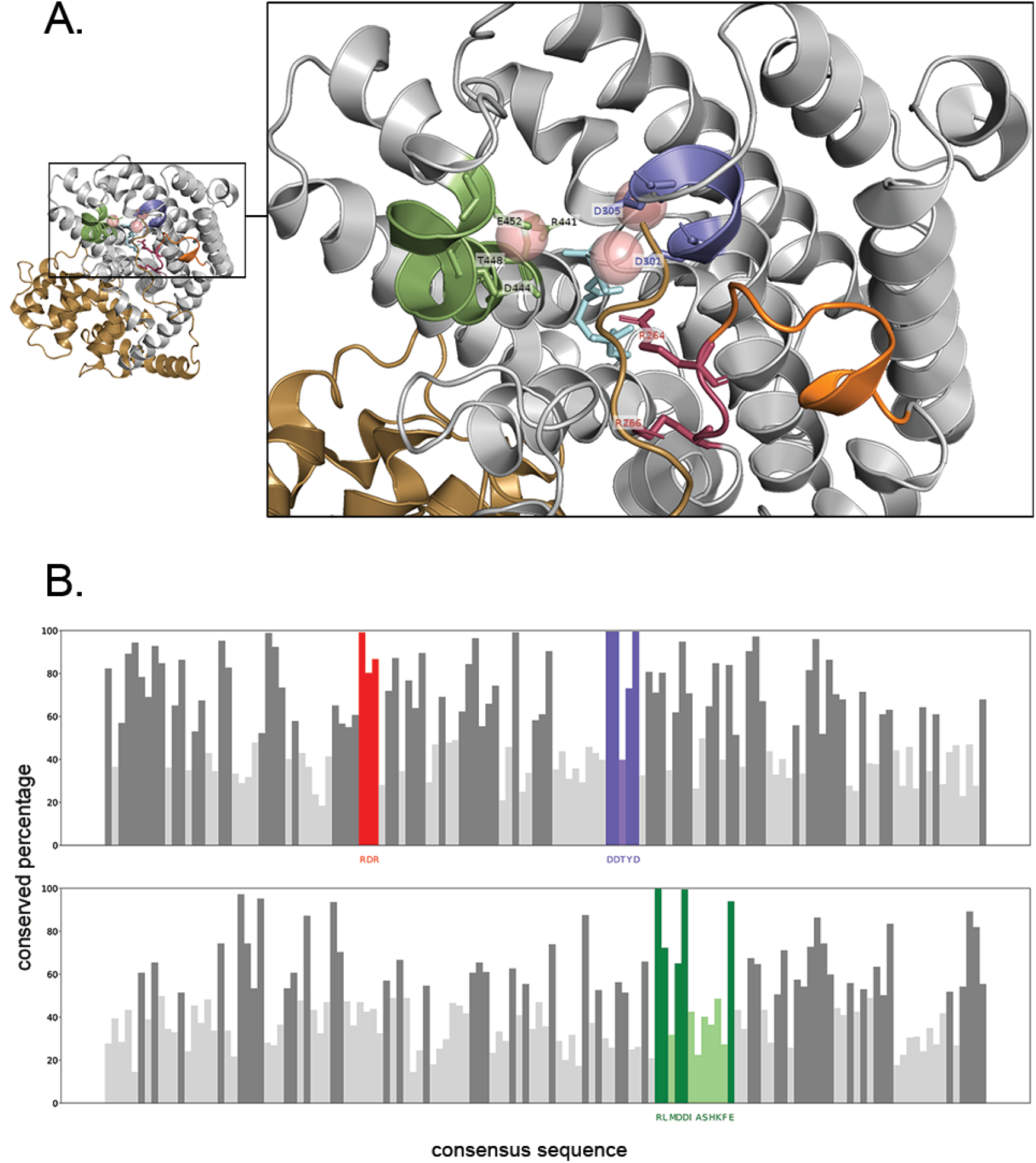
A. Known TPS motifs - RxR (red), DDxxD (purple) and NSE/DTE (green) shown on the structure of tobacco 5-epi-aristolochene synthase (PDB ID: 5EAT). The C-terminal domain is in gray while the N-terminal domain is in brown. Pink spheres represent *Mg*^+2^ ions. A substrate analog, farnesyl hydroxyphosphonate (FHP) is in blue. The A-C loop is coloured in orange. The two conserved Arginines in the <u>R</u>x<u>R</u> motif are shown along with the metal-binding residues in the DDxxD (<u>D</u>Dxx<u>(D,E)</u>) and NSE/DTE motifs (<u>R</u>xx<u>(N,D)</u>Dxx<u>(S,T,G)</u>xxx<u>E</u>). The Arginine in the expanded NSE/DTE motif is also shown, and is found to be very conserved in spermatophyte TPSs. B. The same motifs shown on a schematic of the alignment of all spermatophyte C-terminal sub-sequences from the database. Each bar represents the percentage conservation of the consensus amino acid in the corresponding position of the alignment. Lighter colored bars represent positions where the consensus amino acid is <50% conserved.

#### 2.4.1. Aspartate-rich DDxxD motif conserved in plant STSs

The most conserved motif of TPSs is the metal binding aspartate-rich motif found both in plant and microbial TPSs as well as in isoprenyl diphosphate synthases. Numerous studies performed on this motif, both site-directed mutagenesis and X-ray crystallography analysis, show that it is involved in binding the divalent metal ions in the active site entrance [26]. The canonical form of the motif, **D**Dxx(**D,E**), where bold-faced residues indicate those proposed to bind *Mg*^+2^ or *Mn*^+2^, is found in 247 of the 249 spermatophyte enzymes. Of the remaining two, one is a (+)-germacrene-D synthase from *Solidago canadensis* with an Asn replacing the first Asp [27]. The other is a bicyclogermacrene synthase from *Matricaria chamomilla* with an Asn replacing the second Asp [28]. These examples indicate that either one of the first two Aspartates may be sufficient for maintaining catalytic activity.

#### 2.4.2. Expanded NSE/DTE *motif found in most sequences*

The opposite site of the active site entry is also involved in metal-binding, due to the presence of a second, less-defined motif, termed the NSE/DTE motif [29]. An early form of this motif, as detailed by Christianson *et al*.[29] had a consensus of (L,V)(V,L,A)(**N,D**)D(L,I,V)x(**S,T**)xxx**E**, where the residues in bold coordinate *Mg*^+2^ ions. However, searching for a motif with this consensus only captured 38 of the 249 spermatophyte sequences in our database, indicating that it may be too restrictive given the current knowledge of sequences. When only the metal-binding portion of the motif is considered, the consensus sequence (**N,D**)Dxx(**S,T,G**)xxx**E** covers 219 spermatophyte sequences in the database. The possibility of Gly in the second metal-binding position is justified by Peters *et al*.[30], with the proposal that Gly may allow a water molecule to substitute for the hydroxyl group of Ser/Thr. Some TPSs however, are known to have a second, catalytically active, aspartate rich motif instead of the NSE/DTE motif [31, 32, 33] with the same consensus as the first, **D**Dxx(**D,E**). This occurs in 20 sequences. Table 2 shows the distribution of the sequences over the different versions of the second motif.

**Table 2.**
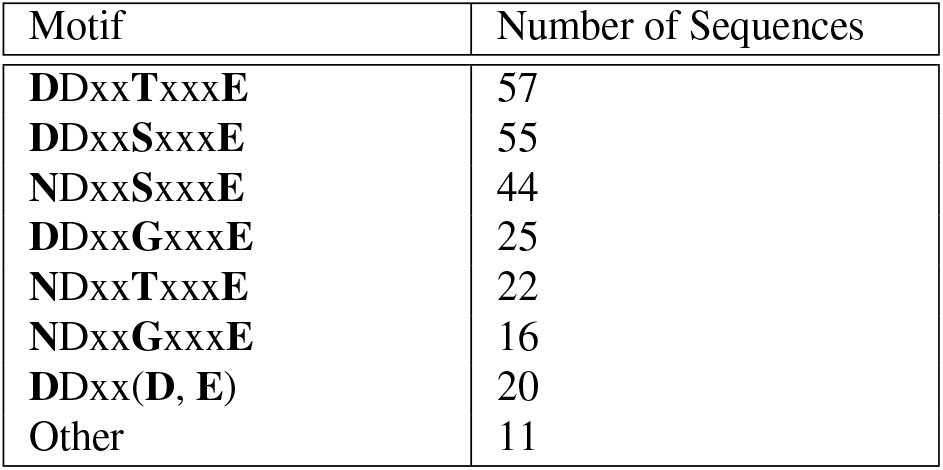
Division of the different versions of the second metal-binding motif among characterized spermatophyte STS sequences. Sequences with motifs not covered by either motif consensus sequence (**N,D**)Dxx(**S,T,G**)xxx**E** or **D**Dxx(**D,E**) are classified as “Other”.

| Motif | Number of Sequences |
| --- | --- |
| DDxxTxxx**E** | 57 |
| DDxxSxxx**E** | 55 |
| NDxxSxxx**E** | 44 |
| DDxxGxxx**E** | 25 |
| NDxxTxxx**E** | 22 |
| NDxxGxxx**E** | 16 |
| DDxx(**D, E**) | 20 |
| Other | 11 |

A highly conserved Arg is found 3 residues upstream of all versions of the NSE/DTE motif or second aspartate-rich motif, in all of the spermatophyte sequences in the database. All 6278 uncharacterized spermatophyte TPS sequences also have an arginine in this position. Hence, an extended form of the motif may be more relevant for spermatophyte STSs, with the consensus Rxx(**N,D**)Dxx(**S,T,G**)xxx**E** or Rxx**D**Dxx(**D,E**).

#### 2.4.3. RxR motif not conserved in nerolidol synthases

The RxR motif is found about 35 amino acids upstream of the DDxxD motif, located on a flexible loop in the structure, termed the A-C loop. This loop has been shown to become ordered upon FPP binding [25]. The two Arg residues in the motif were proposed to be involved in the complexation of diphosphate after ionization of the substrate, thereby preventing nucleophilic attack on any of the carbocationic intermediates [25]. 215 of the 249 spermatophyte plant sequences have the canonical RxR motif while 18 of the remaining have an altered RxQ motif in the same region. Interestingly, these 18 enzymes all catalyze the formation of nerolidol, an acyclic sesquiterpene. This indicates that RxQ may be unable to capture diphosphate to the same extent as RxR, causing a premature quenching of an intermediate carbocation by water before cyclization has occurred [5].

### 2.5. Comparing residues involved in product specificity across sequences

Many studies have addressed the importance of specific residues located in the active site of TPSs via mutational analyses. Some of the best characterized TPSs derive from *Artemisia annua*, which is the source of many medicinal terpenes. Some of the STSs from *A. annua* have served as examples to identify residues involved in critical steps in the cyclization cascade. In this section three examples of *A. annua* STSs are described, for which residues involved in product specificity were experimentally investigated. We use these as a case-study to illustrate how the large set of characterized STSs that we make available can potentially be used to guide such experimental investigations. These examples are:

1. Salmon *et al*. (2015) tested a wide library of mutants for the (*E*)-*β*-farnesene synthase (UniProt: Q9FXY7) from *A. annua*, an STS catalyzing the formation of an acyclic product. They discovered that a single substitution, Tyr402Leu, confers to the synthase a cyclase activity, resulting in zingiberene and *β*-bisabolene as the most abundant products [34]. Both these sesquiterpenes derive from cation 5. In sequences catalyzing the formation of 10,1 and 11,1 cyclized products (cations 1, 2, 3 and 4), this position is highly conserved (88-100%) in the database as a Tyr, and Leu does not occur. However, STSs producing cation 5 and those producing acyclic products have relatively lower conservation in this position (70% Tyr and 53% Phe respectively) and Leu is found 14% of the time in cation 5. Thus conservation patterns in this position are indicative of the corresponding residue’s contribution to product specificity.
2. In another study, Li *et al*. (2013) studied the effect of mutations on the cyclization reaction of the bisabolol synthase from *A. annua* (UniProt: M4HZ33) [35]. A possible reaction mechanism involves formation of a nerolidyl cation, followed by the formation of cation 5 by a 1,6 ring closure, and deprotonation to produce the final product bisabolol [36]. The authors identified a mutation that interfered with this 1,6 ring closure and showed that the substitution Leu399Thr changed the product specificity, to *γ*-humulene, derived from cation 2, a 11,1 cyclization of the farnesyl cation [35]. Interestingly, a Leu at this position is quite rare; it is present in only four sequences in the database, all four of which belong to the group of sequences producing cation 5. Instead, this position is highly conserved (>95%) as either a Ser or a Thr in the database.
3. Amorpha-4,11-diene is a bicyclic sesquiterpene produced from the 6,1-cyclized bisabolyl cation, cation 5 in Figure 1. Li *et al*. (2016) did a mutational analysis of the amorpha-4,11-diene synthase from *A. annua* (UniProt: Q9AR04), and showed that the residue Thr296 can cause a loss of cyclization activity when mutated [37]. This residue is 82% conserved as either a Ser or a Thr in cyclic STSs. Importantly, in acyclic STSs the most common amino acid is a Tyr, with a conservation of 38%. Acyclic STSs even have amino acids such as Gln, Gly and Ile in this position, never seen in the cyclic STSs in the database. The variability and low conservation score indicates that changing this position in cyclic STSs away from a Ser or Thr could result in the formation of acyclic products, as shown by Li *et al*. [37].

In summary, analysis of these *A. annua* examples of residues involved in the first cyclization step in STSs indicates that conservation patterns across all the annotated enzymes are consistent with the functional roles of these residues. This suggests it would be possible to obtain residues potentially involved in product specificity from this database. Such a data-driven approach is in contrast to how these mutational studies have traditionally been guided, i.e. by comparison of two or three sequences from the same or related species. Therefore, a potential application of our database is in guiding site-directed mutagenesis studies in a way which avoids species bias and hence may reveal additional residues involved in product specificity. One such residue position obtained by studying conservation patterns has been discussed above in Section 2.4.3, namely the second arginine in the RxR motif. This position was found to be glutamine in most nerolidol synthases, something not seen in any of the cyclic synthases. Mutating this residue in cyclic synthases and monitoring for acyclic products, and vice versa, could confirm the residue’s role in the cyclization of sesquiterpene products.

## 3. Conclusion

We compiled a manually curated set of experimentally characterized plant STSs along with their major products. This database is the largest centralized resource of annotated plant STSs to date and allows for thorough sequence-based analysis of these diverse enzymes. The enzymes in the database are grouped according to the carbocationic origin and cyclization of their major product. Such a division alleviates the task of functional analysis and comparison between the enzymes. Using the database we were able to extend and find variants of existing STS motifs. In addition, residues from previous mutational studies, when mapped onto the enzymes in the database, were found to have detectable conservation patterns that differed from group to group. Such properties of residues can be extrapolated and used to guide further mutational studies. The database as a whole helps to understand the current state of STS sequence space characterization, and provides a starting point for future efforts to predict product specificity.

## 4. Experimental

### 4.1. Literature search for characterized STSs

To find potentially characterized STSs, an HMM search was performed using hmmer (version 3.1b2) [38] on the UniProt database [39] using the HMM of the C-terminal domain of TPSs from Pfam [20] (Pfam ID: PF03936). Protein sequences with a hit having an E-value < 10^−10^ and a total protein length between 350 and 650 residues were selected. The Uniprot IDs of these sequences were then linked to Pubmed IDs, either directly through programmatic access of Uniprot if the Pubmed ID was present, or through a programmatic text search of the title and authors given in Uniprot, using the Pubmed API [40]. The Pubmed articles thus obtained were searched manually for evidence of experimental characterization of sesquiterpenes through in-vivo or in-vitro GC-MS studies, and the corresponding Uniprot IDs were collected.

For each UniProt ID found, the major product described in the corresponding paper was stored. Minor products with GC-MS peaks at least quarter the height of the major product peak were stored as well.

### 4.2. Measuring chemical similarities

The diagram of the sesquiterpene grouping scheme was made using ChemDoodle (version 9) [41]. The InChI strings for 165 sesquiterpenes were obtained from PubChem [42] using the python wrapper for the PubChem REST API [43], PubChemPy (version 1.0.4). To measure the similarity between different sesquiterpenes, rdkit (Release 2017.09.3) was used [44]. A circular chemical fingerprint, called the Morgan fingerprint, with a radius of 2 angstroms, as explained by Rogers et al [22], was obtained for each sesquiterpene. The similarity between every pair of fingerprints was then calculated using Dice similarity [21]. The distance was simply given as 1 − *similarity*. The distance matrix of all sesquiterpenes was then used to create a multi-dimensional scaling (MDS) plot using the Python scikit-learn library (version 0.19.1) [45], and then plotted using matplotlib (version 2.1.2) [46].

### 4.3. Aligning sequences

For characterized spermatophyte plant STS sequences, the C-terminal catalytically active portion of the enzyme was found with a hmmer HMM search (version 3.1b2) [38] using the TPS C-terminal Pfam domain (Pfam ID: PF03936). These were then aligned using Clustal Omega (version 1.2.4) [47], with all heuristic features off and the Pfam domain as a guide for alignment.

For some of the nonseed plant STS sequences however, a Pfam domain search returned <200 residues instead of the usual 250-270. Aligning the full nonseed sequences using the spermatophyte C-terminal sub-sequence alignment as a profile showed the position of the C-terminal portion for these sequences, so this was used to extract the required sub-sequences for nonseed plants. An alignment consisting of both seed and nonseed characterized C-terminal subsequences was constructed using Clustal Omega with the same parameters as above.

### 4.4. Phylogenetic tree construction

A phylogenetic tree was built and visualized for the characterized spermatophyte and nonseed plant enzymes in the database using the ete toolkit (version 3.1.1) [48]. The previously explained alignment of all C-terminal subsequences was used, with columns having >50% gaps removed using trimAL [49]. The best protein model from JTT, WAG, VT, LG and MtREV was chosen using ProtTest [50], and finally a RaxML maximum likelihood tree was built [51].

### 4.5. Finding mono-, di-, and uncharacterized TPSs

Characterized plant mono- and diterpene synthases were obtained from SwissProt [11] using a C-terminal TPS Pfam domain hmmer (verson 3.1b2) [38] HMM search followed by collecting the sequences from plant species for which the catalytic activity was mentioned. These were not manually checked.

Uncharacterized TPS C-terminal sub-sequences were then obtained from plant species in TremBl [11], Ensembl Plants (release 38) [52], and the 1000 Plants Transcriptome Project [53] again using a Pfam domain search. Only those sequences where the search returned a sub-sequence having both DDxx(D,E) and (N,D)Dxx(S,T,G)xxxE or two DDxx(D,E) motifs within it, and whose sub-sequence length was within two standard deviations of the mean C-terminal sub-sequence length of characterized STS enzymes were retained. In both sets, sequences from nonseed plant species were discarded.

### 4.6. Measuring sequence similarities

A distance matrix of all spermatophyte TPS C-terminal sub-sequences: characterized mono-, di- and sesquiterpene synthases as well as uncharacterized enzymes, was constructed using the pairwise sequence k-tuple measure described by Wilbur *et al*. [54], implemented in Clustal Omega (version 1.2.4) [47]. This distance matrix was then used to construct an MDS plot using scikit-learn (version 0.19.1) [45] and plotted using matplotlib (version 2.1.2) [46]. A cluster-map of sequence identities between characterized STS enzymes was made using the distance matrix of just these enzymes and complete hierarchical clustering using scipy (version 1.0.0) [55] and seaborn (version 0.8.1).

### 4.7. Visualizing an STS structure

The 5EAT tobacco 5-epi-aristolochene synthase structure from the Protein Data Bank (PDB) [56] was used to visualize known TPS motifs, along with *Mg*^+2^ ions and farnesyl hydroxyphosphonate (FHP) substrate analog. Visualization was done in Pymol 2.1 [57].

## Acknowledgements

This work is part of the research programme Novel Enzymes for Flavour and Fragrance with project number TTW 15043 which is financed by the Netherlands Organisation for Scientific Research (NWO).

